# Structural basis for the HMGCR interaction with UBIAD1 mutants causing Schnyder corneal dystrophy

**DOI:** 10.1101/2020.06.29.177683

**Authors:** Fengbo Zhou, Jayne S. Weiss, Weikai Li

## Abstract

Schnyder corneal dystrophy (SCD) is an autosomal dominant disease characterized by abnormal deposition of cholesterol and lipid in the cornea. The molecular mechanism underlying this process, which involves the interaction between UBAID1 and HMGCR, remains unclear. Here we investigate these events with *in silico* approaches. We built the homology models of UBIAD1 and HMGCR based on the existing crystal and cryo-EM structures. The UBIAD1 and HMGCR models are docked and their binding interactions are interrogated by MD simulation. We find that the transmembrane helices of UBIAD1 bind to sterol sensing domain of HMGCR. Upon binding of the GGPP substrate, UBIAD1 shows lower structural flexibility in the TM regions binding to HMGCR. The N102S and G177R mutations disrupts GGPP binding, thereby lowering the binding affinity of HMGCR. Overall, our modeling suggests that SCD mutations in UBIAD1 or lower GGPP concentration increase the structural flexibility of UBIAD1, thereby facilitating its association with HMGCR.

## Introduction

Schnyder corneal dystrophy (SCD) is an autosomal dominant disease that affects the front portion of the eye called the cornea. SCD is characterized by an abnormal amount of cholesterol and lipid deposits in the usually transparent cornea, leading to corneal opacification and loss of vision (Weiss, 1992; Weiss, 2007). The relentless deposition of corneal lipid results in a predictable pattern of corneal opacification with age. While laser treatment can ablate crystals that may deposit in the visual axis and impair vision, progressive corneal lipid deposition continues. There is no medical treatment to prevent or slow the accumulation of lipid in the SCD cornea. Currently, only surgical excision and replacement of the cornea can be offered once the vision has already been impaired from corneal opacification. Thus, understanding the pathogenesis of SCD is critical in order to develop therapeutic interventions that could treat or prevent the disease.

SCD is caused by mutations in UbiA prenyltransferase domain containing 1 (*Ubiad1*) (Orr *et al*, 2007; Weiss *et al*, 2007), which encodes an intramembrane prenyltransferase that synthesizes vitamin K2. The UBIAD1 enzyme catalyzes the condensation reaction of geranylgeranyl pyrophosphate (GGPP) with menadione, which is originated from the dietary vitamin K1 (Nakagawa *et al*, 2010). Many SCD mutations, such as Asn102Ser and Gly177Arg, impair the catalytic activity of this enzyme (Nickerson *et al*, 2013; Nickerson *et al*, 2010).

The mechanism of cholesterol and lipid accumulation in SCD is yet to be clarified, despite the knowledge that UBIAD1 interacts with several proteins involved in the cholesterol and lipid metabolism (Fredericks *et al*, 2013; Nickerson *et al.*, 2013). One possibility is insufficient removal of lipids from the cornea. The C-terminal portion of apolipoprotein E, a constituent of HDL is known to interact with *UBIAD1.* This interaction is associated with solubilization of cholesterol and its removal from cells. Thus, it is possible that *UBAID1* mutations result in reduced cellular cholesterol removal and increased accumulation of corneal HDL (Weiss *et al.*, 2007). An alternative explanation is that excess cholesterol production could be linked to alternation in the mevalonate pathway. This pathway generates both sterols, such as cholesterol, and nonsterol isoprenoids. such as GGPP, which is the UBIAD1 substrate. The common precursor of these compounds, mevalonate, is generated by HMG-CoA reductase (HMGCR), the rate-limiting enzyme of the mevalonate pathway (Sever *et al*, 2003). HMGCR is regulated by multiple feedback mechanisms that may involve UBIAD1. Through regulating the mevalonate pathway, SCD mutations in UBIAD1 can cause cholesterol accumulation.

Indeed, SCD mutations in UBIAD1 have been found to interfere with the regulation pathway of HMGCR through endoplasmic reticulum (ER) associated protein degradation (ERAD) (Schumacher *et al*, 2015; Schumacher *et al*, 2016). The ERAD of HMGCR is accelerated by sterol, but sterol also triggers the HMGCR binding to UBIAD1 to inhibit this ERAD process, thereby constituting a feedback loop. GGPP promotes the release of HMGCR from UBIAD1, which is normally cycled between ER and Golgi. Conversely, UBIAD1 becomes retained in the ER with either decreasing level of GGPP or the SCD mutations. The trapped UBIAD1 in turn blocks the HMGCR degradation; the increased HMGCR activity that promotes the mevalonate synthesis explains the cholesterol accumulation associated with SCD mutations.

The molecular mechanisms underlying the regulated association and dissociation of UBAID1 and HMGCR remain unclear. Here we investigate these events with *in silico* approaches. We built the homology model of UBIAD1 by using crystal structures of its archaeal homologs (Cheng & Li, 2014a; Huang *et al*, 2014). For HMGCR, we combined the crystal structure of its soluble domain with the model of transmembrane domain, which is based partly on the known structures of the cholesterol-sensing domain (Gong *et al*, 2016; Li *et al*, 2016; Qi *et al*, 2019). Subsequently, the UBIAD1 and HMGCR models are docked and their binding interactions are interrogated by MD simulation. We find that the transmembrane helices of UBIAD1 bind to sterol sensing domain of HMGCR. Upon binding of the GGPP substrate, UBIAD1 shows lower structural flexibility in the TM regions binding to HMGCR. The N102S and G177R mutations disrupts GGPP binding, thereby lowering the binding affinity of HMGCR. Overall, our modeling suggests that SCD mutations in UBIAD1 or lower GGPP concentration increase the structural flexibility of UBIAD1, thereby facilitating its association with HMGCR.

## Results

### Homology model of UBIAD1 structure

In order to generate a UBIAD1 homology model, we first compared the sequences of human UBIAD1 and two archaeal homologs from *Aeropyrum pernix* (ApUbiA) and *Archaeoglobus fulgidus* (AfUbiA) (Figure 1A). These homologs are selected because their crystal structures have been determined (Cheng & Li, 2014a; Huang *et al.*, 2014). The sequence identity and similarity are 19.0% and 32.6% between human UBIAD1 and ApUbiA, 21.2 % and 36.8% between human UBIAD1 and AfUbiA, and 24.3% and 41.8% between ApUbiA and AfUbiA, respectively. Such moderate sequence similarity is not uncommon for homologous membrane proteins, whose transmembrane regions appear to have relative high tolerance to sequence variations. Despite the moderate sequence similarities, the overall structures of the two archaeal homologs are essentially the same, suggesting that the UBIAD1 structure can be reliably modeled. The sequence alignment, however, shows that these two homologs do not contain the first 49 residues of human UBIAD1 (Figure 1A). TMHMM prediction of UBIAD1 shows that these N-terminal 49 residues are not in the transmembrane region. UBIAD1 contains nine transmembrane helices (TMs) as the archaeal homologs do, and the boundary of the TMs from TMHMM prediction are almost the same as the structural observation for the homologs. Therefore, we only modeled the rest of the UBIAD1 structure to ensure reliability.

**Figure 1.**
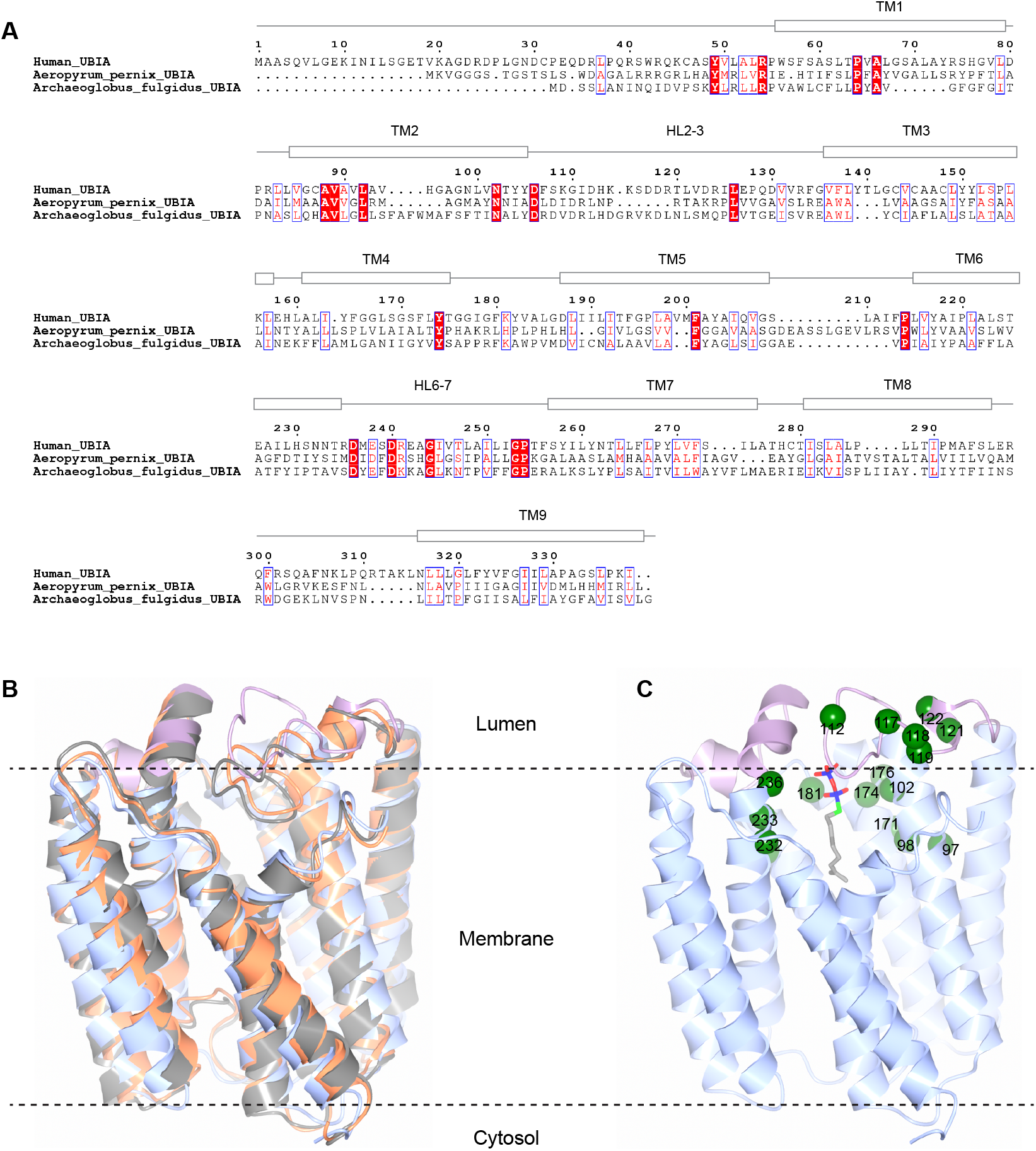
The UBIAD1 homology model. **A,** Sequence alignment of human UBIAD1, ApUbiA and AfUbiA. UBIAD1 alignment between full length human UBIAD1 (HsUbiA) and archaeal UBIAD1 homologs (Ap_UbiA, and Af_UbiA). UBIAD1 is conserved across the species in residues 60-338 of human UBIAD1. **B,** Superposition of the UBAD1 model and the crystal structures of ApUbiA and AfUbiA. **C.** Homology model of UBAD1. The green spheres indicate location of SCD mutations.

We modeled the UBIAD1 structure by using I-TASSER with one-to-one threading of either of the homolog structures. Moreover, we generated the homology model of UBIAD1 using automatic search of the sequence database. These three homology models show essentially the same structure (Figure 1B). Superimposing this UBIAD1 model onto the crystal structures of ApUbiA and AfUbiA show a low RMSD of 1.9 Å and 1.5 Å, respectively. Taken together, the homology modeling based on the crystal structures provide a relatively reliable model of UBIAD1 for subsequent investigations (Figure 1C). In this model, the two DXXXD motifs and Asn102 are at the active site, where the substrate GGPP is expected to be bound.

### Modeling of the sterol-sensing domain of HMGCR

HMGCR is comprised of an N-terminal transmembrane domain and a C-terminal soluble region (489-871) carrying the HMG-CoA reductase activity. Crystal structure of this catalytic domain has been determined (Istvan *et al*, 2000), but that of the transmembrane domain is unknown. In absence of this structural information, our early attempts using automatic homology modeling was insufficient to generate a meaningful model of the full-length HMGCR. Thus, the transmembrane domain of HMGCR needs to be properly modeled first. Previous studies of membrane topology show that HMGCR have eight transmembrane helices (TMs) (Olender & Simon, 1992). Notably, the TM 2-6 (residues 88–218) of HMGCR is a conserved sterol-sensing domain (SSD). Biologically, sterol is important in promoting the HMGCR binding to UBIAD1. Thus, modeling the SSD is an important step towards understanding the regulated UBIAD1-HMGCR interactions.

For SSD-containing proteins, structures known to date are those of the human Niemann–Pick C1 protein (NPC1) (Gong *et al.*, 2016; Li *et al.*, 2016) and the human hedgehog receptor, PTCH1 (Qi *et al.*, 2019). The SSDs of NPC1 (TM3-TM7; residues 620-771) and HMGCR (TM 2-6; 61–204 AA) share a 21.5% sequence identity and 43.7% similarity (Figure 2A). Such moderate levels of sequence similarity are also found between the SSD of PTCH1 (TM2-TM6; AA 285-471) and NPC1, and between the SSD of PTCH1 and HMGCR. TMHMM prediction of the boundary of these transmembrane regions in HMGCR are almost the same as in the NPC1 and PTCH1 structures (Figure 2A). Likewise, these structures show that the transmembrane regions of NPC1 and PTCH1 are essentially the same, with a RMSD of 2.1 Å (Figure 2B). Based on these structural templates, we generated a homology model of HMGCR sterol-sensing domain (Figure 2C). The modeling used I-TASSER with one-to-one threading based on either of the homolog structures. These two homology models show essentially the same structure and superposition of the models gives a RMSD of 2.1 Å (Figure 2C). Taken together, structure of the HMGCR SSD can be reasonably modeled (Figure 2C).

**Figure 2.**
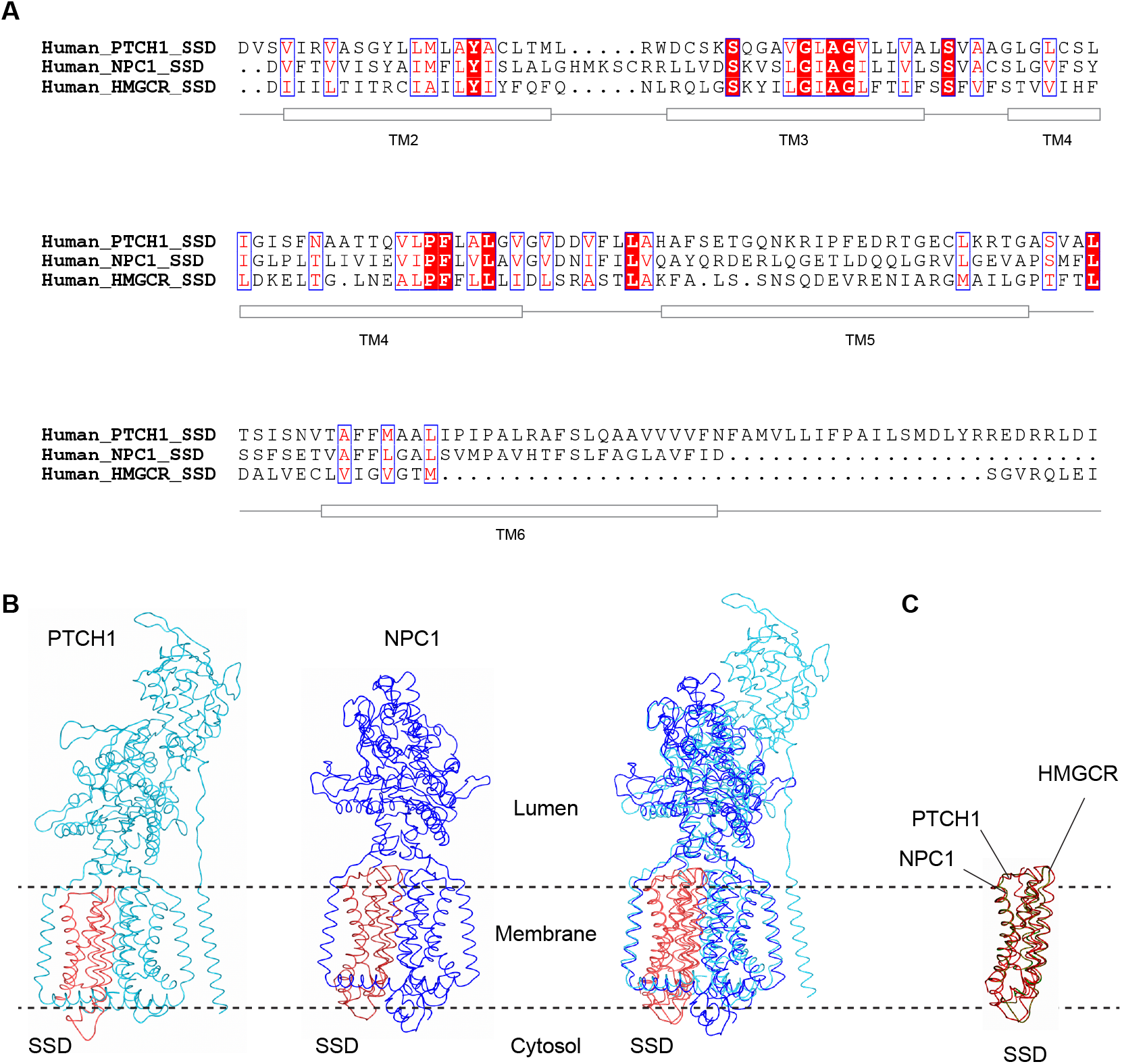
Homology model of the HMGCR sterol-sensing domain. **A,** Sequence alignment of the sterol-sensing domain of human HMGCR, NPC1 and PTCH1. **B,** Sterol-sensing domain in NPC1 (orange) and PTCH1 (red). **C.** Superposition of the model of HMGCR sterol-sensing domain with those of NPC1 and PTCH1.

### Modeling of the full-length HMGCR

With the SSD of HMGCR modeled, the rest of its transmembrane regions flanking the SSD (1-87 and 219-440) remain with unknown structures. Therefore, we generated the homology models of each of these individual regions by using automatic search of the database. The models show that the region 1-60 contains a TM in between two loops. The region 219-440 contains a hairpin of two TMs connected by a loop. Together, the automatic generation of these three TMs, plus the five TMs of SSD, is well consistent with topology experiments showing that the HMGCR contains eight TMs (Olender & Simon, 1992). Therefore, we combined these TMs together to generate the initial model of the transmembrane domain (Figure 3A). This combined model was energy minimized. Subsequently, we performed MD simulation to assess the stability of this model. We find that the model maintains a folded eight-TM structure over the 100 ns simulation time (Figure 3B).

**Figure 3.**
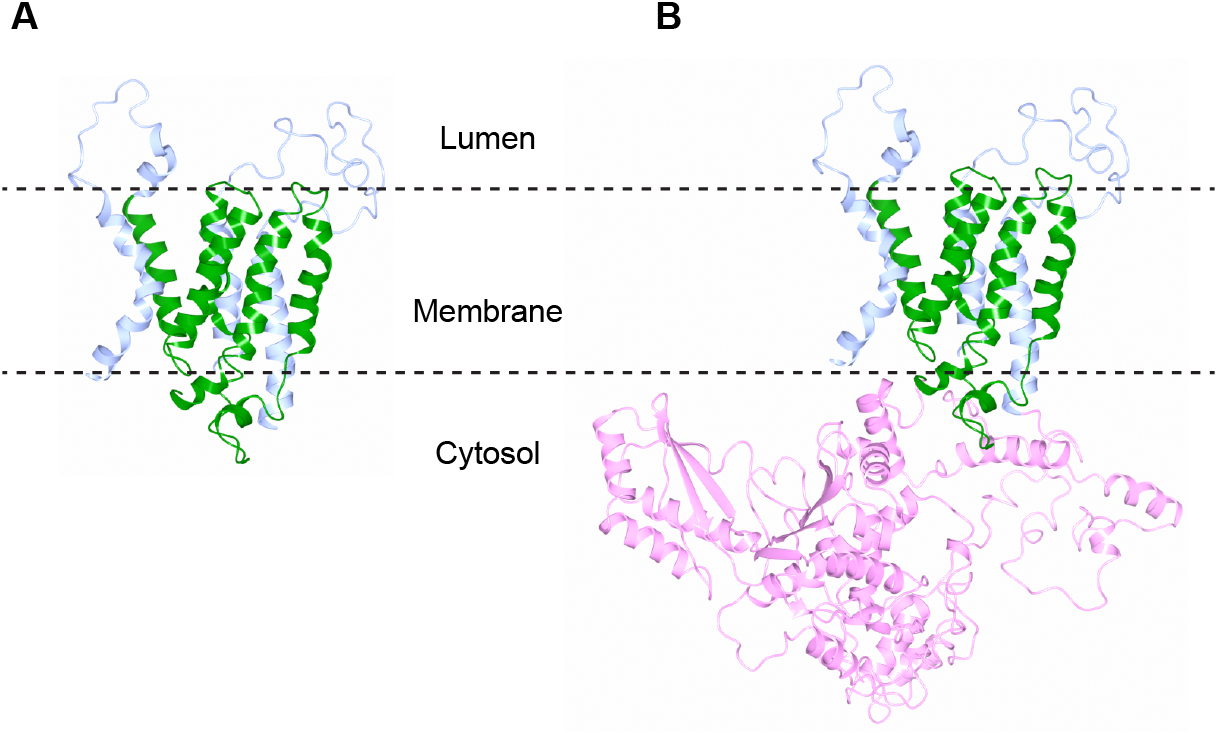
Model of HMGCR. **A,** Homology model of HMGCR transmembrane domain. The sterol-sensing domain is shown in green and additional TMs in blue. **B,** Model of the full length HMGCR.

To generate the model of full-length HMGCR, we fused the crystal structure of the HMGCR catalytic domain with the modeled eight-TM structure (Figure 3C). For the connecting region (352-440) between the two domains, homology modeling shows that this region is mostly unstructured, consistent with the secondary structure prediction. Subsequently, we performed MD simulation of this full-length model (Figure 3D). During the 100 ns simulation, we observed some level of interactions between the transmembrane and catalytic domains. Despite the limitation of our modeling, the potential interactions of these two HMGCR domains suggest a regulation mechanism of the reductase activity through sterol binding (Luo *et al*, 2020).

### Modeling of the HMGCR-UBIAD1 complex

To understand the interaction between HMGCR with UBIAD1, we performed the docking of UBIAD1 homology model with the models of full-length HMGCR. Because most membrane proteins interact through their transmembrane domains, we only selected docking models that the transmembrane domains of HMGCR and UBIAD1 contact with each other. Moreover, the docking model needs to be consistent with the membrane orientation of HMGCR and UBIAD1; the catalytic domain of HMGCR faces the cytosol (Olender & Simon, 1992) whereas the UBIADI active site faces the ER lumen (Li, 2016). Therefore, we selected docking models that these two catalytic sites face opposite sides of the membrane plane. We further selected the model with the highest score and lowest free energy. This model shows that the TM1-5 of UBIAD1 contacts the SSD of HMGCR (Figure 4), consistent with the notion that cholesterol binding affects the interaction between UBIAD1 and HMGCR (Schumacher *et al.*, 2015; Schumacher *et al.*, 2016). Subsequently, we inserted this model of the HMGCR-UBIAD1 complex into the POPC lipid bilayer and performed energy minimization of the entire system.

**Figure 4.**
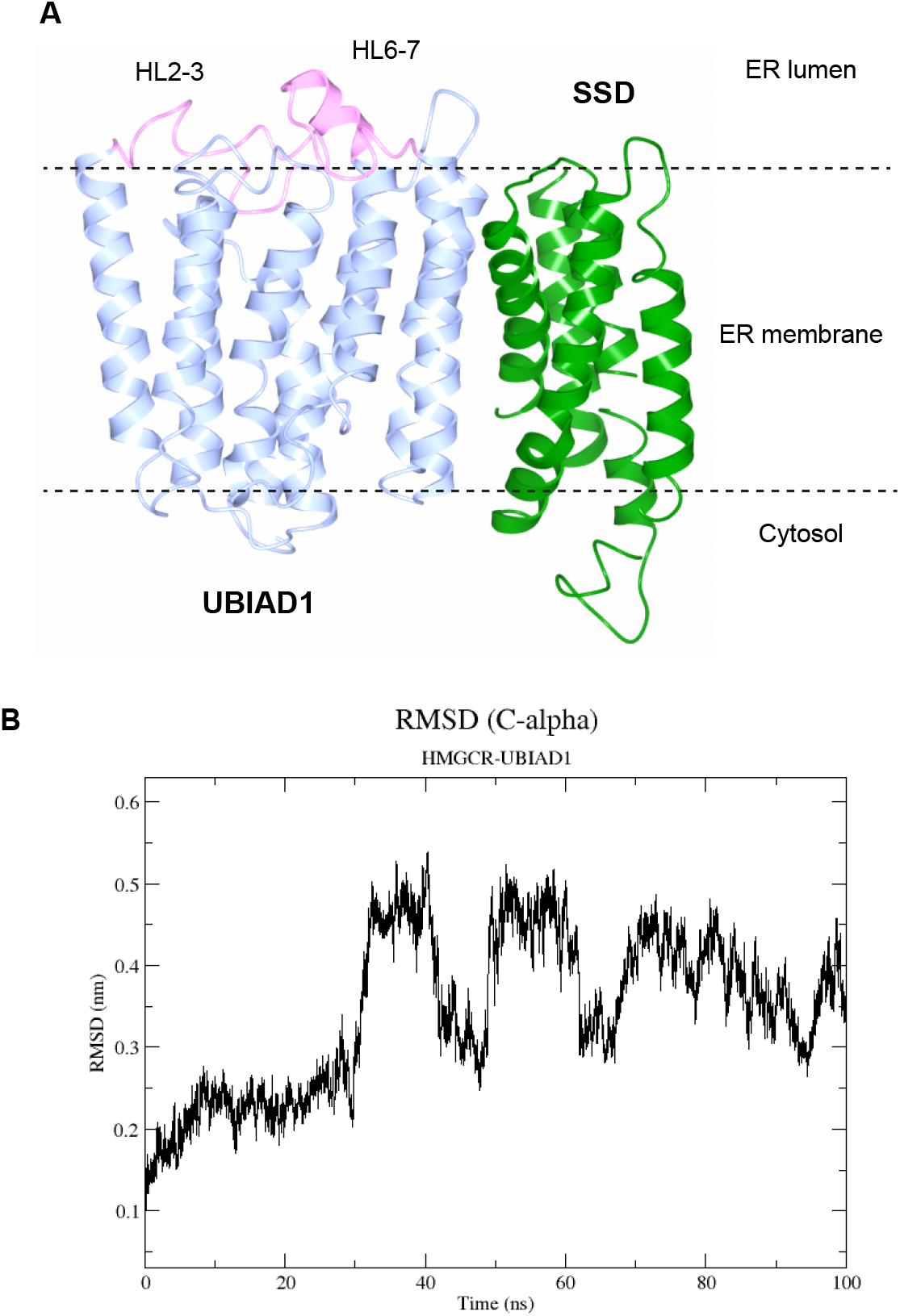
Model and MD simulation of the HMGCR-UBIAD1 complex. **A,** Model of the HMGCR-UBIAD1 complex. Only the sterol-sensing domain is shown. **B,** RMSD of MD simulation of the HMGCR-UBIAD1 complex.

### Molecular dynamics simulation of HMGCR-UBIAD1 interaction

We performed molecular dynamics simulation for the HMGCR-UBIAD1 complex model. During 100-ns simulation, orientation of the HMGCR-UBIAD1 complex is not tilted and its overall conformation remain unchanged in the POPC lipid bilayer (Figure 4B). The over position of UBIAD1 relative to HMGCR is also unchanged, indicating that the binding interaction between HMGCR and UBIAD1 complex is stably formed in the MD model. The binding interface between UBIAD1 and HMGCR involves multiple interactions between the sterol-sensing domain and the transmembrane domain of UBIAD1. We did not observe any interaction between the HMGCR catalytic domain and UBIAD1, probably due to the small exposed regions of this integral membrane protein. Instead, the sterol-sensing domain of HMGCR provide the major UBIAD1-interaction site throughout the simulation that samples possible movements between the two molecules.

### Modeling and molecular dynamics simulation of UBIAD1 with GGPP

The UBIAD1 substrate, GGPP, promotes the dissociation of UBIAD1 from HMGCR. To understand how GGPP binding affects the dissociation, we generated a UBIAD1-GGPP binding complex. Using the UBIAD1 homology model, we inserted a GGPP molecule to make a model with the bound substrate. The pyrophosphate and isoprenyl chain of GGPP are docked into the UBIAD1 model, positioned and orientated in the same way as GPP found in the crystal structure of the archaeal homolog. This is because the pyrophosphate binding sites, with the two DXXXD motif and surrounding arginine residues, are conserved between the archaeal UbiA homologs and the UBIAD1. Subsequently, the UBIAD1-GGPP complex was embedded into POPC lipid bilayer. After energy minimization, we performed 20 ns MD simulation of this complex system. The conformation of UBIAD1 with GGPP bound remains stable during the simulation time (Figure 5).

**Figure 5.**
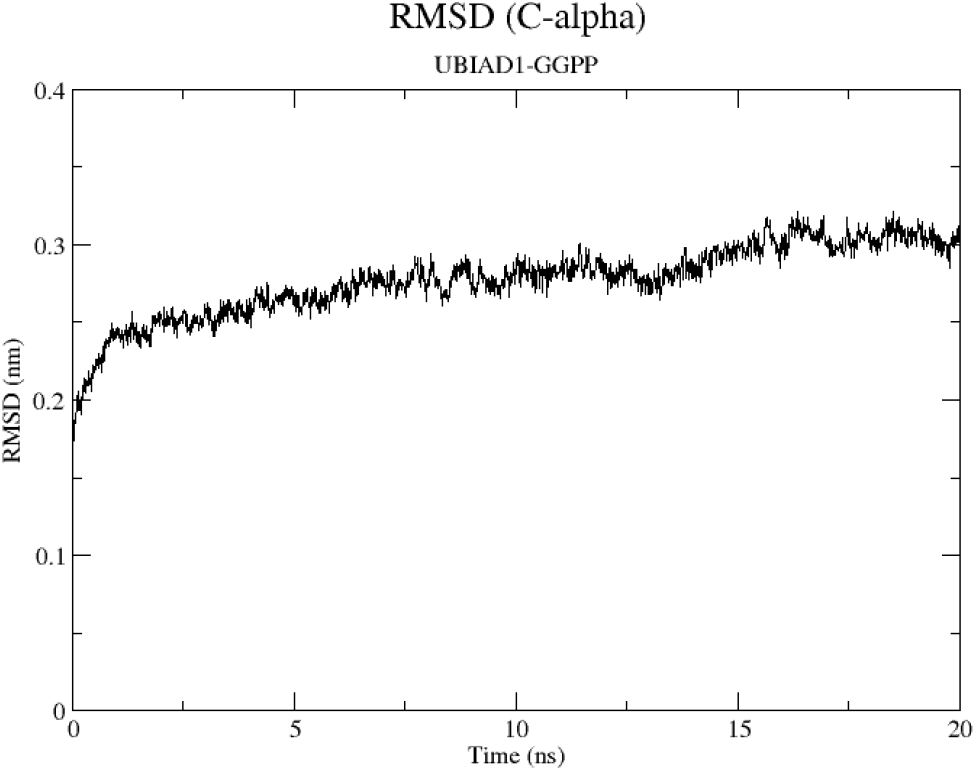
RMSD of MD simulation of the UBIAD1-GGPP complex.

## Discussion

The stability of the GGPP bound structure of UBIAD1 is consistent with previous biochemical analyses showing that GGPP binding lower the structural flexibility of the UbiA homologs (Cheng & Li, 2014b; Huang *et al.*, 2014). GGPP binding to UBIAD1 may have a similar effect of tightening its structure. Conversely, release of GGPP from UBIAD1 renders its conformational flexibility, which is probably required for UBIAD1 to adopt an alternative conformation that is required for HMGCR association. This model explains regulation of HMGCR degradation through the GGPP binding and release in UBIAD1.

Most SCD mutations are clustered around the active site (Figure 1C). The most prominent SCD mutation, N102S, is directly involved in the catalysis. It is therefore likely that these SCD mutations act by abolishing the GGPP binding. This generates the same effect as the release of this substrate. The constitutive absence of the substrate may promote the UBIAD1 association with HMGCR. As explained above, formation of this stable complex prevents the HMGCR degradation by ERAD. Consequently, the excess activity of HMGCR causes cholesterol accumulation though enhancing the mevalonate pathway. In sum, the SCD mutants of UBIAD1 lead to cholesterol accumulation in the cornea due to impaired GGPP binding.

The binding of UBIAD1 and HMGCR are likely to interact at the sterol-sensing region of HMGCR, supported by the fact that the binding between UBIAD1 and HMGCR is stimulated by sterols. Moreover, a yeast to hybrid (Y2H) liquid culture assay showed that overlapping HMGCR clones were isolated with a minimal prey fragment containing HMGCR residues 1–285, a region that included the 5TM sterol sensing region in HMGCR.

The SCD disease has been thought to be a local metabolic abnormality of the cornea, because progressive corneal lipid deposition and opacification occurs independent of serum cholesterol values (Lisch *et al*, 1986). While two thirds of affected SCD individuals also have hypercholesterolemia; systemic hypercholesterolemia is also noted in many unaffected members of the SCD pedigree (Weiss, 2007). Thus, future studies are required to elucidate why the disregulated UBIAD1-HMGCR interaction leading to cholesterol accumulation occurs mainly in the cornea. Overall, understading the SCD pathology at the molecular level is a key step towards the development of new therapeutic strategies for SCD.

## Methods

The homology models were generated using I-TASSER (Roy *et al*, 2010) and PHYRE2 (Kelley *et al*, 2015). The docking of UBIAD1 and HMGCR homology models used the Z-DOCK online program (Pierce *et al*, 2014). Among the ten predictions of the binding from the docking results, we selected three models which showed close binding between the SSD of HMGCR and UBIAD1. We submitted the three prediction models for energy minimization, and a subsequent 1-ns MD production run to test and to stabilize the docking model (Fig. 2). The final stabilized UBIAD1-HMGCR (5-TM) model was yielded from the top prediction that that was not only correct in terms of topology of UBIAD1 and HMGCR, but also had the face SCD mutation sites contacting the SSD of HMGCR.

Molecular dynamics simulation was performed with GROMACS MD simulation package (Van Der Spoel *et al*, 2005) with CHARMM36m force field (Huang *et al*, 2017). For MD simulation, the system was first energy minimized with a steepest descent algorithm with a 0.01-nm step size and a 1.2-nm neighbor-list cutoff distance for Coulomb and Van der Waals interactions until the maximum force on any atom in the system fell below 1,000 kJ/mol/nm. The system was then equilibrated for 0.25 ns with a 0.001-fs time step and a Berendsen thermostat to hold the system temperature at 303.15 K. A Verlet cutoff scheme was used for the neighbor list. Particle mesh Ewald was used for the Coulomb interactions. Parrinello–Rahman barostat and velocity-rescaling thermostat was used for the MD simulation.

## References

Cheng W, Li W (2014a) Structural Insights into Ubiquinone Biosynthesis in Membranes. Science (New York, NY) 343: 878–881

Cheng W, Li W (2014b) Structural insights into ubiquinone biosynthesis in membranes. Science 343: 878–881

Fredericks WJ, McGarvey T, Wang H, Zheng Y, Fredericks NJ, Yin H, Wang L-P, Hsiao W, Lee R, Weiss JS et al (2013) The TERE1 (UBIAD1) bladder tumor suppressor protein interacts with mitochondrial TBL2: regulation of trans-membrane potential, oxidative stress and SXR signaling to the nucleus. Journal of cellular biochemistry 114: 2170–2187

Gong X, Qian H, Zhou X, Wu J, Wan T, Cao P, Huang W, Zhao X, Wang X, Wang P et al (2016) Structural insights into the Niemann-Pick C1 (NPC1)-mediated cholesterol transfer and ebola infection. Cell 165: 1467–1478

Huang H, Levin EJ, Liu S, Bai Y, Lockless SW, Zhou M (2014) Structure of a membrane-embedded prenyltransferase homologous to UBIAD1. PLoS biology 12: e1001911–e1001911

Huang J, Rauscher S, Nawrocki G, Ran T, Feig M, de Groot BL, Grubmuller H, MacKerell AD, Jr. (2017) CHARMM36m: an improved force field for folded and intrinsically disordered proteins. Nat Methods 14: 71–73

Istvan ES, Palnitkar M, Buchanan SK, Deisenhofer J (2000) Crystal structure of the catalytic portion of human HMG-CoA reductase: insights into regulation of activity and catalysis. EMBO J 19: 819–830

Kelley LA, Mezulis S, Yates CM, Wass MN, Sternberg MJE (2015) The Phyre2 web portal for protein modeling, prediction and analysis. Nature Protocols 10: 845–858

Li W (2016) Bringing Bioactive Compounds into Membranes: The UbiA Superfamily of Intramembrane Aromatic Prenyltransferases. Trends in Biochemical Sciences 41: 356–370

Li X, Wang J, Coutavas E, Shi H, Hao Q, Blobel G (2016) Structure of human Niemann–Pick C1 protein. Proceedings of the National Academy of Sciences 113: 8212–8217

Lisch W, Weidle EG, Lisch C, Rice T, Beck E, Utermann G (1986) Schnyder’s dystrophy. Progression and metabolism. Ophthalmic Paediatr Genet 7: 45–56

Luo J, Yang H, Song BL (2020) Mechanisms and regulation of cholesterol homeostasis. Nat Rev Mol Cell Biol 21: 225–245

Nakagawa K, Hirota Y, Sawada N, Yuge N, Watanabe M, Uchino Y, Okuda N, Shimomura Y, Suhara Y, Okano T (2010) Identification of UBIAD1 as a novel human menaquinone-4 biosynthetic enzyme. Nature 468: 117–121

Nickerson ML, Bosley AD, Weiss JS, Kostiha BN, Hirota Y, Brandt W, Esposito D, Kinoshita S, Wessjohann L, Morham SG et al (2013) The UBIAD1 prenyltransferase links menaquinone-4 [corrected] synthesis to cholesterol metabolic enzymes. Human mutation 34: 317–329

Nickerson ML, Kostiha BN, Brandt W, Fredericks W, Xu K-P, Yu F-S, Gold B, Chodosh J, Goldberg M, Lu DW et al (2010) UBIAD1 mutation alters a mitochondrial prenyltransferase to cause Schnyder corneal dystrophy. PloS one 5: e10760–e10760

Olender EH, Simon RD (1992) The intracellular targeting and membrane topology of 3-hydroxy-3-methylglutaryl-CoA reductase. J Biol Chem 267: 4223–4235

Orr A, Dubé M-P, Marcadier J, Jiang H, Federico A, George S, Seamone C, Andrews D, Dubord P, Holland S et al (2007) Mutations in the UBIAD1 gene, encoding a potential prenyltransferase, are causal for Schnyder crystalline corneal dystrophy. PloS one 2: e685–e685

Pierce BG, Wiehe K, Hwang H, Kim BH, Vreven T, Weng Z (2014) ZDOCK server: interactive docking prediction of protein-protein complexes and symmetric multimers. Bioinformatics 30: 1771–1773

Qi C, Di Minin G, Vercellino I, Wutz A, Korkhov VM (2019) Structural basis of sterol recognition by human hedgehog receptor PTCH1. Sci Adv 5: eaaw6490

Roy A, Kucukural A, Zhang Y (2010) I-TASSER: a unified platform for automated protein structure and function prediction. Nature protocols 5: 725–738

Schumacher MM, Elsabrouty R, Seemann J, Jo Y, DeBose-Boyd Ra (2015) The prenyltransferase UBIAD1 is the target of geranylgeraniol in degradation of HMG CoA reductase. eLife 4: 1–21

Schumacher MM, Jun D-J, Jo Y, Seemann J, DeBose-Boyd RA (2016) Geranylgeranyl-regulated transport of the prenyltransferase UBIAD1 between membranes of the ER and Golgi. Journal of Lipid Research 57: 1286–1299

Sever N, Yang T, Brown MS, Goldstein JL, DeBose-Boyd RA (2003) Accelerated degradation of HMG CoA reductase mediated by binding of insig-1 to its sterol-sensing domain. Molecular Cell 11: 25–33

Van Der Spoel D, Lindahl E, Hess B, Groenhof G, Mark AE, Berendsen HJC (2005) GROMACS: Fast, flexible, and free. Journal of Computational Chemistry 26: 1701–1718

Weiss JS, Schnyder’s dystrophy of the cornea: A Swede-Finn connection.

Weiss JS (1992) Schnyder’s dystrophy of the cornea. A Swede-Finn connection. Cornea 11: 93–101

Weiss JS (2007) Visual morbidity in thirty-four families with Schnyder crystalline corneal dystrophy (an American Ophthalmological Society thesis). Transactions of the American Ophthalmological Society 105: 616–648

Weiss JS, Kruth HS, Kuivaniemi H, Tromp G, White PS, Winters RS, Lisch W, Henn W, Denninger E, Krause M et al (2007) Mutations in the UBIAD1 gene on chromosome short arm 1, region 36, cause Schnyder crystalline corneal dystrophy. Investigative ophthalmology & visual science 48: 5007–5012

